# Lipid network of plasma extracellular vesicles reveals sex-based differences in the lipidomic profile from patients with alcohol use disorder

**DOI:** 10.1101/2023.09.12.557479

**Authors:** Carla Perpiñá-Clérigues, Susana Mellado, Cristina Galiana-Roselló, María Fernández-Regueras, Miguel Marcos, Francisco García-García, María Pascual

## Abstract

Alcohol use disorder (AUD) is one of the most common psychiatric disorders, and the consumption of this substance is considered one of the main causes of preventable deaths worldwide. Lipids play a crucial functional role in cell membranes, however little is known about the role of lipids containing extracellular vesicles (EVs), as regulatory molecules and biomarkers. In this study, a highly sensitive lipidomic strategy is employed to characterize plasma EV lipid species from individuals with AUD, to evaluate differential functional roles and enzymatic activity networks to improve the knowledge of lipid metabolism in the alcohol consumption. Plasma EV lipids from female and male patients with AUD and healthy individuals were analyzed to obtain lipid differential abundance, as well as biological interpretation of LINEX^2^ lipidomics data, evaluating enzymatic dysregulation through an enrichment algorithm. Our results showed for the first time that females with AUD exhibit greater substrate-product changes in LPC and PC lipids, as well as phospholipases and acyltransferases activity, potentially linked to cancer progression and neuroinflammation. Conversely, males with AUD showed dysregulation in Cer and SM lipid, involving sphingomyelinases, sphingomyelin phosphodiesterase, and sphingomyelin synthase, which could be related with hepatotoxicity. Notably, females with AUD showed LION-terms associated with “positive intrinsic curvature”, while males exhibited “negative intrinsic curvature, contributing to vesicle fusion processes. These methodological developments allow a better understanding of lipid metabolism and its regulatory mechanisms, which contributes not only to identify novel lipid targets, but also the discovery of sex-specific clinical biomarkers in the AUD.

## Introduction

Alcohol use disorder (AUD) is a chronic disease characterized by unhealthy alcohol use and several neurobiological features that can include positive reinforcement, compulsive search for alcohol and negative emotional state when alcohol is not used [1]. It consists of a constellation of symptoms, including withdrawal, tolerance and craving, among others. AUD is a major public health issue, the prevalence of which has been increasing at an alarming rate. Indeed, alcohol use is estimated to cause approximately 3 million deaths globally per year, constituting a major factor for morbimortality [2]. Alcohol has several health consequences, such as alcohol-associated liver disease, hepatocellular carcinoma, non-liver neoplasms, physical injury, cardiac disease, and psychiatric disorders. Alcohol misuse significantly affects workforce productivity, with elevated direct and indirect economic costs. Unfortunately, a major proportion of people affected by alcohol are in the most productive years of their lives [2].

Extracellular vesicles (EVs) are diverse, nanoscale membrane vesicles actively released by cells. These microvesicles are increasingly recognized as important vehicles of intercellular communication and circulating biomarkers for disease diagnoses and prognosis [3]. A range of studies has demonstrated the role of EVs in physiological processes and pathological conditions, such as inflammation, cancer, and neurodegenerative diseases [4]. While recent research has provided extensive information concerning the role of DNA, RNA and protein content of EVs in biological processes, little is known about the role of EV lipids. We have recently demonstrated that binge-like ethanol drinking induces a differential enrichment of EV lipid species in human female adolescents compared to males. Furthermore, these lipid species participate in EV formation, release, and uptake, and inflammatory immune response [5].

Lipids play a crucial role in cell membranes as well as participate in various cellular functions. Understanding how lipid changes caused by pathological conditions, environmental factors, or treatments, impact cellular processes is vital, and provides new insights into potential disease mechanisms [6]. Dedicated computational tools are essential for this aim [7]. Mass spectrometry-based lipidomics, combined with computational analysis, is a powerful tool for identifying and quantifying lipids in cells, tissues, or bodily fluids [8]. Although recent reports are focused on lipid composition and abundance, thousands of lipids interact via many pathways and networks. Evaluating and understanding changes in these networks in response to cellular environment alterations, in association with the development of a disease, are crucial to deciphering cell metabolism and related molecular mechanisms [9]. In this sense, the development of new bioinformatic tools, such as lipid network explorer (LINEX^2^) which combines lipid class and fatty acid metabolism, provides comprehensive networks for computational analysis and interpretation of lipidomics data [7]. These methodological developments allow a better understanding of lipid metabolism and its regulatory mechanisms, which contributes not only to identify novel drug targets, but also the discovery of clinical biomarkers [10, 11].

Taking into account the novel approach based on bioinformatic analysis and the critical roles of EV lipids as biomarkers, we employed a highly sensitive lipidomic strategy to characterize EV lipid species isolated from plasma of male and female individuals with AUD, and evaluate the differential functional roles and enzymatic activity networks of the EV lipids to improve the role of lipid metabolism in the alcohol consumption. We demonstrated sex-based differences in the EVs lipid composition induced by alcohol consumption, impacting species and class-level abundance, as well as lipid metabolic networks. Furthermore, these lipids were found to be related to EVs biogenesis and/or to the inflammatory and neurodegenerative responses. All data and results generated have been made openly available on a web-based platform (http://bioinfo.cipf.es/sal-chronics).

## Materials and methods

### Human subjects

We included a total of 11 patients (6 males and 5 females) with AUD according to DSM-5 criteria that were referred to the Alcoholism Unit of the University Hospital of Salamanca. The median age of the subjects was 47.83 and 40.00 for males and females with AUD, respectively. As previously described [12], all patients included in this group were actively drinking (≥ 90 g of ethanol/day) until entering the study. All patients had normal prothrombin time, hemoglobin concentration and albumin serum levels as well as negative hepatitis B surface antigen and antibodies to hepatitis C virus. They did not have other chronic or acute conditions that could alter the results of the study or were polydrug abusers. Advanced liver disease was excluded based on clinical, analytical and ultrasonographic studies: individuals displaying physical stigmata of chronic liver disease (e.g., cutaneous signs, hepatosplenomegaly, gynecomastia, testicular atrophy, and/or muscle wasting), with liver ultrasonographic findings other than steatosis or with increased liver transaminases > 2-3 times the reference limits were excluded. In addition, 12 healthy volunteers (6 males and 6 females) were also analyzed, who reported to drink < 15 g of ethanol/day and they all showed normal liver function tests and standard hematological and biochemical tests (see Table S1). The median age of the healthy males and females was 45.50 and 39.50, respectively. Before entering the study, all individuals gave their informed consent to participate, and the study was approved by the Ethics Committee of the University Hospital of Salamanca.

Heparin-anticoagulated peripheral blood samples were obtained from the patients and healthy patients, between 9:00 and 10:00 AM, under fasting conditions. Plasma samples were snap frozen in liquid nitrogen and stored at −80 °C until further use. Samples were processed for biochemical tests and EV isolation. Figure 1A summarizes the human experimental groups.

**Figure 1.**
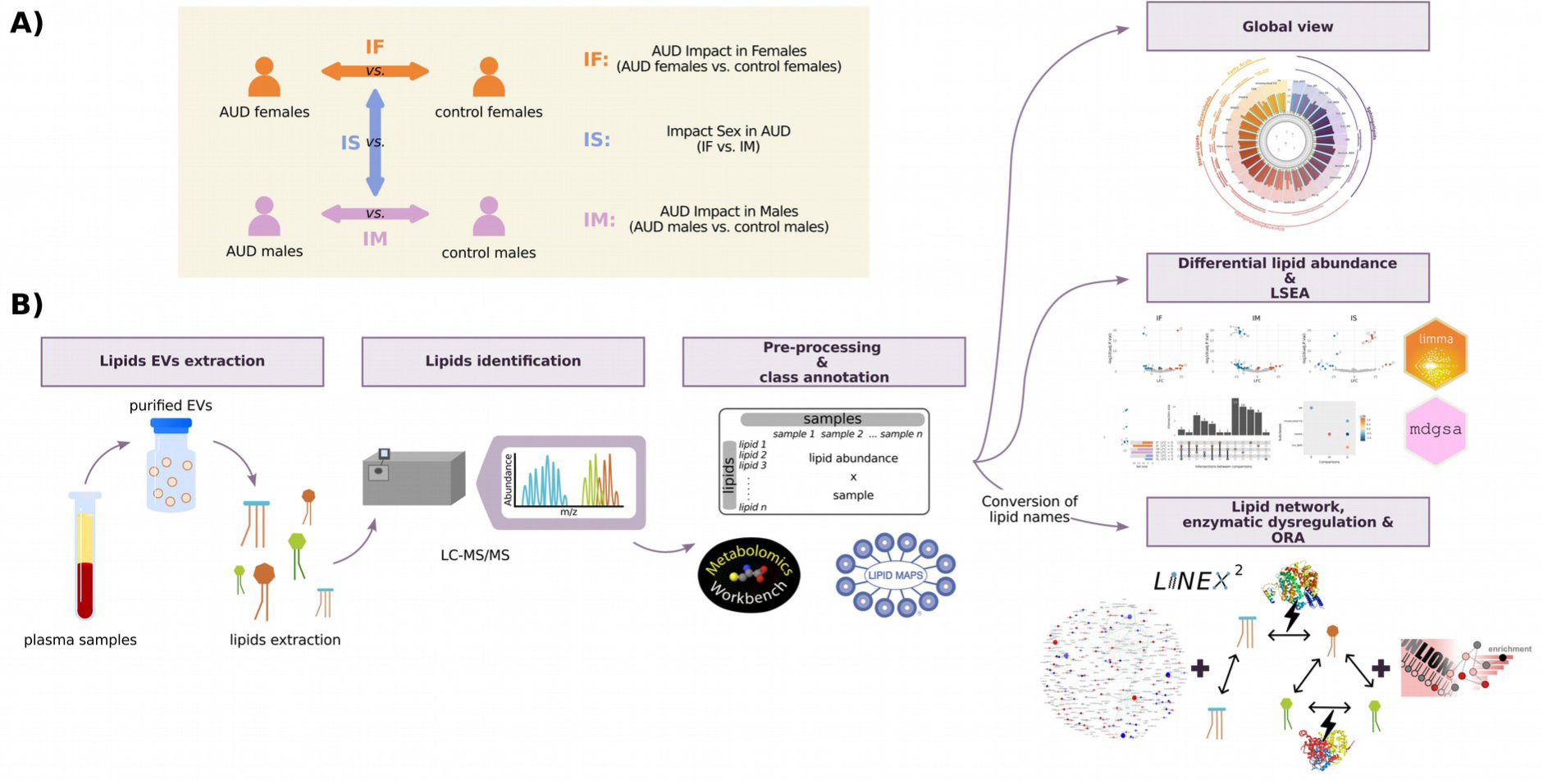
Lipidomic workflow, describing the subjects analyzed and comparisons performed in the different analyses. **(A)** Experimental groups and comparisons. **(B)** After isolating EVs from human plasma, lipids were extracted for quantification and identification through LC-MS/MS. Following data normalization and lipid class annotation, exploratory and differential analyses were conducted to assess lipid abundance. A class enrichment analysis was also performed. Additionally, utilizing the LINEX2 platform, we obtained a reaction global network, the heaviest connected subgraph, and a target lipids list for further enrichment analysis using LION-web.

### EV isolation from human plasma

Plasma EVs were isolated by a total exosome isolation kit (catalog number 4484450, Invitrogen, USA), following the manufacturer’s instructions. 250 μL of initial plasma was used to isolate EVs, and these were collected and frozen at −80°C until processing.

### EVs characterization by transmission electron microscopy and nanoparticles tracking analysis

Freshly isolated EVs were fixed with 2% paraformaldehyde and were prepared as previously described [13]. Preparations were examined under a transmission FEI Tecnai G2 Spirit electron microscope (FEI Europe, Eindhoven, The Netherlands) with a digital camera Morada (Olympus Soft Image Solutions GmbH, Münster, Germany). In addition, an analysis of the absolute size range and concentration of microvesicles was performed using NanoSight NS300 Malvern (NanoSight Ltd., Minton Park, UK), as previously described [13]. Figure 2 reports EVs characterization by electron microscopy and nanoparticles tracking analysis.

**Figure 2.**
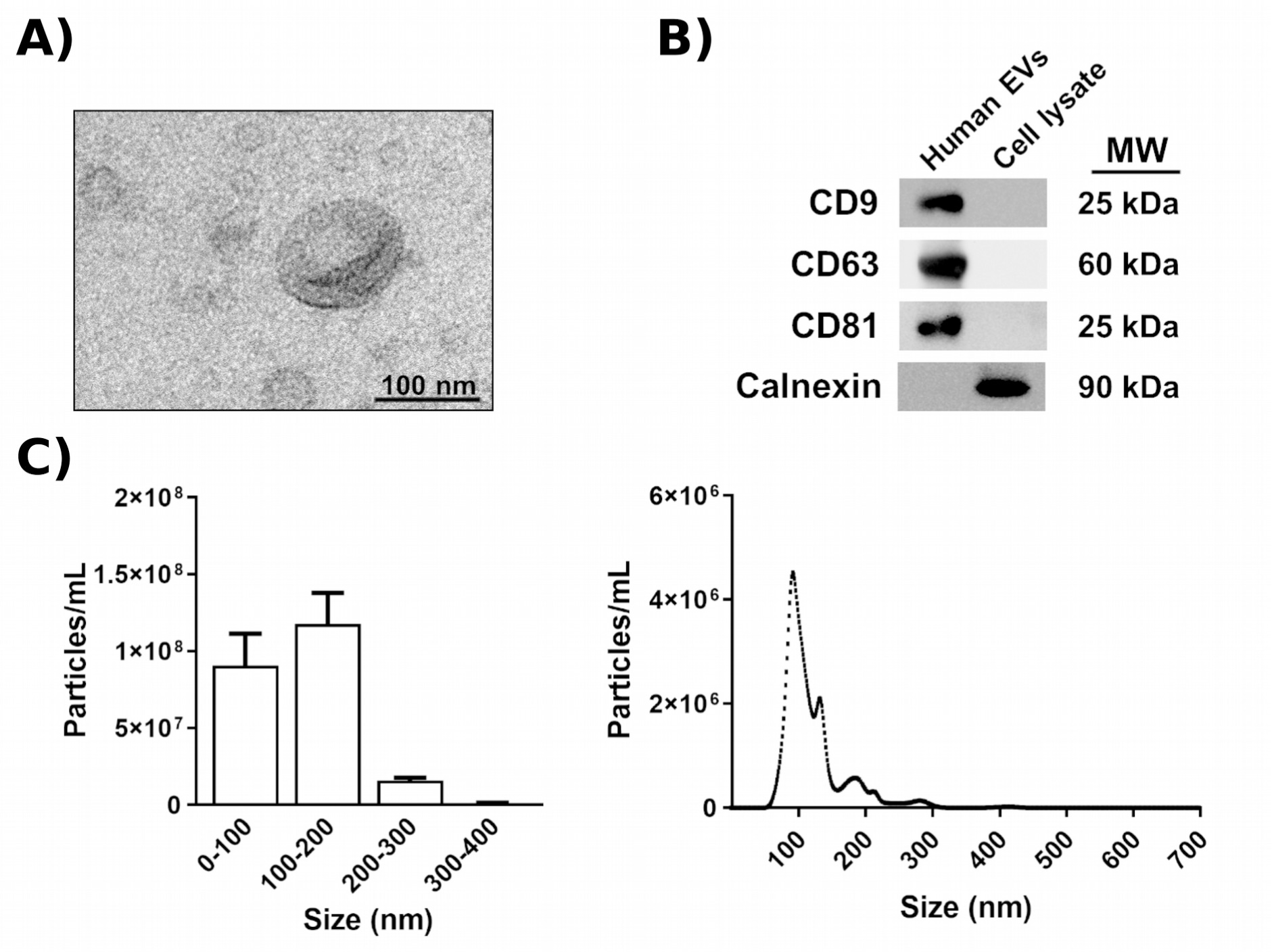
Characterization of plasma EVs. **(A)** Electron microscopy image of human plasma EVs. **(B)** Analysis of the protein expression of EV markers (CD9, CD63, and CD81) in EVs and cell lysates. Calnexin expression was used to discount cytosolic protein contamination in EV samples. Cell lysates from astroglial cells were used as a positive control for calnexin expression. A representative immunoblot for each protein is shown. **(C)** Measurement of human EV size distribution and concentration by nanoparticle tracking analysis. A high peak ranging between 100 and 200 nm is shown, which includes the size range of EVs.

### Western blot analysis

The Western blot technique was performed in the plasma EVs for their characterization (Fig. 2), as described elsewhere [14]. The primary antibodies used were: anti-CD9, anti-CD63, anti-CD81 and anti-calnexin (Santa Cruz Biotechnology, USA). Membranes were washed, incubated with the corresponding HRP-conjugated secondary antibodies and developed using the ECL system (ECL Plus; Thermo Fisher Scientific). Figure S1 in the Supplementary Material includes the whole membrane of each protein expression.

### Lipid extraction

Lipids were extracted from equal amounts of plasma-derived EVs (0.2 mL/sample), using a modified Folch extraction procedure. The last phase containing the lipids was transferred to fresh tubes, dry vacuumed with nitrogen and lipids were stored at −80°C until further analysis. Dried samples were then resuspended with isopropanol for different LC/MS acquisition methods (positive and negative ion modes).

### LC-MS/MS analysis

In fully automated Q-TOF acquisition mode, a pooled human lipid extract representing the 36 samples (4 conditions × 9 replicates) was acquired by iterative MS/MS. Detailed experimental methods for chromatography and autoMS/MS mass spectrometry were followed as described [15, 16] with minor modifications. Briefly, sample separation was performed using an Agilent 1290 Infinity LC system coupled to the 6550 Accurate-Mass QTOF (Agilent Technologies, Santa Clara, CA, USA) with electrospray interface (Jet Stream Technology) operating in positive-ion mode (3500 V) or negative-ion mode (3000 V) and high sensitivity mode. The optimal conditions for the electrospray interface were gas temperature of 200 °C, drying gas 12L/min, nebulizer 50 psi, sheath gas temperature of 300 °C, and sheath gas flow 12 L/min. Lipids were separated on an Infinity Lab Poroshell 120 EC-C18 column (3.0 ×100 mm, 2.7 μm) (Agilent, Santa Clara, CA, USA). Under optimized conditions, the mobile phase consisted of solvent A (10 mM ammonium acetate, 0.2 mM ammonium fluoride in 9:1 water/methanol) and solvent B (10mM ammonium acetate, 0.2 mM ammonium fluoride in 2:3:5 acetonitrile/methanol/isopropanol) using the following gradient: 0 min 70%B, 1 min 70%B, 3.50 min 86%B, 10 min 86%B, 11 min 100%B, 17 min 100%B operating at 50 °C and a constant flow rate of 0.6 mL/min. Injection volume was 5 µL for positive and negative modes.

The Agilent Mass Hunter Workstation Software was employed for the data acquisition. LC/MS Data Acquisition B.10.1 (Build 10.1.48) was operated in auto MS/MS, and the three most intense ions (charge states, 1-2) within 300–1700 m/z mass range (over a threshold of 5000 counts and 0.001%) were selected for MS/MS analysis. The quadrupole was set to a “narrow” resolution (1.3 m/z), and MS/MS spectra (50–1700 m/z) were acquired until 25,000 total counts or an accumulation time limit of 333 ms. To assure the desired mass accuracy of recorded ions, continuous internal calibration was performed during analyses using signals m/z 121.050873 and m/z 922.009798 for positive mode and signals m/z 119.03632 and m/z 980.016375 for negative mode. Additionally, all-ions MS/MS [17] data were acquired on individual samples, with an MS acquisition rate of three spectra/second and four scan segments 0, 10, 20, and 40 eV.

### Lipid annotator database

5 sets of 5 iterative MS/MS data files from pooled human cell extracts were analyzed with Lipid Annotator software 1 as the first step in the lipidomics workflow. This study used a novel software tool (Lipid Annotator) [18] with a combination of Bayesian scoring, a probability density algorithm, and non-negative least-squares fit to search a theoretical lipid library (modified LipidBlast) developed by Kind et al. [19, 20] to annotate the MS/MS spectra.

Agilent MassHunter Lipid Annotator Version 1.0 was used for all other data analyses. Default method parameters were used, except only [M+H]+ and [M+NH4] + precursors were considered for positive ion mode analysis, and only [M-H]– and [M+HAc-H]– precursors were considered for negative ion mode analysis. Agilent MassHunter Personal Compound Database and Library (PCDL) Manager Version B.08 SP1 was used to manage and edit the exported annotations.

### Lipid identification

The lipid Personal Compound Database and Library (PCDL) databases created were used for Batch Targeted Feature Extraction in Agilent Mass Hunter Qualitative version 10.0 on the respective batches of 36 all-ions MS/MS data files. The provided “Profinder - Lipids.m” method was adapted in Mass Hunter Qualitative software with modifications previously described by Sartain, M. et al., 2020 [16]. Data were analyzed using the Find by Formula (FbF) algorithm in MassHunter Qualitative Analysis. This approach uses a modified version of the FbF algorithm, which supports the all-ions MS/MS technique. Mass peaks in the low energy channel are first searched against the PCDL created for compounds with the same m/z values, and then a set of putative identifications is automatically compiled. For this list, the fragment ions in the MS/MS spectra from the PCDL are compared to the ions detected in the high-energy channel to confirm the presence of the correct fragments. The precursors and productions are extracted as ion chromatograms and evaluated using a coelution score. The software calculates a number that accounts for abundance, peak shape (symmetry), peak width, and retention time. The resulting compounds were reviewed in the Mass Hunter Qualitative version, and features not qualified were manually removed. Mass Hunter Qualitative results and qualified features were exported as a .cef file.

### Bioinformatic analyses

The strategy applied for this study was based on a transcriptomic analysis workflow. All bioinformatics and statistical analyses were performed using R software v.4.1.2 [21]. Figure 1A illustrates the experimental design and Figure 1B displays the whole lipidomic workflow.

### Data preprocessing

Data preprocessing included filter entities, normalization of abundance lipid matrix, and exploratory analyses. Mass Hunter Qualitative results (.cef file) were imported into Mass Profiler Professional (MPP) (Agilent Technologies) for statistical analysis. Entities were filtered based on their frequency, selecting those consistently present in all replicates of at least 1 experimental group. A percentile shift normalization algorithm (75%) was used, and datasets were baselined to the median of all samples. Normalized data were labeled according to negative and positive ion mode, and all data were consolidated into a single data frame. This step was followed by exploratory analysis using hierarchical clustering, principal component analysis (PCA), and box and whisker plots by samples and lipids to detect abundance patterns between samples and lipids and batch effects anomalous behavior in the data. At this point, anomaly-behaving samples and outliers (values that lie over 1.5 x interquartile range (IQR) below the first quartile (Q1) or above the third quartile (Q3) in the dataset) were excluded for presenting a robust batch effect with a critical impact on differential abundance analysis.

### Differential lipid abundance

Lipid abundance levels between groups were compared using the limma R package [22]. P-values were adjusted using the Benjamini & Hochberg (BH) procedure [23], and significant lipids were considered when the BH-adjusted p-value ≤ 0.05.

### Class enrichment analysis

Class annotation was conducted using the *RefMet* database [24] and compared with the *LIPID MAPS* database [25]. Description of abbreviations is detailed in Table S2. Annotation was followed by ordering lipids according to the p-value and sign of the statistic obtained in the differential lipid abundance. Similar to a Gene Set Enrichment Analysis (GSEA) method, a class enrichment analysis was carried out using Lipid Set Enrichment Analysis (LSEA) implemented in the mdgsa R package [26]. The p-values were corrected for BH, and classes with a BH-adjusted p-value ≤ 0.05 were considered significant.

### Lipid network

The Lipid Network Explorer platform (LINEX^2^, https://exbio.wzw.tum.de/linex/) was used for lipid metabolic network analysis to gain insights into the sex-specific dysregulation of lipid metabolism in AUD patients [7]. For this purpose, single lipid species were considered, either as sum or molecular species, regardless of their retention time and ion mode acquisition. For that reason, prior to conducting the analysis, the lipid nomenclature was checked to ensure that the majority of the lipids in the study were included. This review was carried out using the MetaboAnalyst 5.0 platform [27] and the LipidLynxX Converter tool (http://www.lipidmaps.org/lipidlynxx/converter/) [28]. Additionally, a manual revision was performed on a lipid-by-lipid basis to ensure accuracy. LINEX ^2^ analysis brought forth several results. The global network of lipid species provides both qualitative and quantitative associations between species based on defined reaction types and Spearman’s correlation, respectively. In addition, changes in lipid levels between different experimental conditions can be derived from different statistical metrics. The subgraph with the largest average substrate-product changes was obtained through a lipid network enrichment algorithm, which took enzymatic multispecificity into account and generated hypotheses regarding enzymatic dysregulation. This algorithm consists of a local search approach which generally examines a search space in a greedy manner by iteratively testing local candidate solutions for the one with an optimal objective function. Candidate solutions are generated by applying one of three operations: node insertion, deletion, and substitution to the solution from the last iteration or a randomly selected subgraph in the first iteration. Lastly, LINEX^2^ provided a target lipids list derived from the lipids subgraph, which was utilized for an enrichment analysis using LION-web (http://www.lipidontology.com/) [29]. This enabled a more in-depth examination of the functional significance and potential biological implications of the identified lipid alterations.

### Comparisons

Three comparisons were performed to analyze differential lipid abundance (Fig. 1A): i) the AUD Impact in Females (IF), which compares females with AUD and control females (AUD.Females - Control.Females); ii) the AUD Impact in Males (IM), which compares males with AUD and control males (AUD.Males - Control.Males); iii) Impact of Sex in AUD (IS), which compares IF and IM ((AUD.Females - Control.Females) - (AUD.Males - Control.Males)). Class enrichment analysis was assessed using the same three principal comparisons. LINEX^2^ analysis related to the global network was conducted using IF and IM comparisons, and the subgraph with the largest average substrate-product changes was obtained using the control groups as a reference. The purpose of performing the IF and IM comparisons was to identify the lipids whose abundance was affected by the alcohol consumption separately in each sex. The IS comparison allowed us to identify the lipids whose abundance differed due to sex in the context of AUD.

The statistics used to measure the differential patterns were the logarithm of fold change (LFC) to quantify the effect of differential lipid abundance, and the logarithm of odds ratio (LOR) to measure the enrichment of each functional class. A positive statistical sign indicates a higher mean for the variable in the first element of the comparison, whereas a negative statistical sign indicates a higher mean value for the second element. The IS comparisons focus on finding differences between female and male comparisons. Thus, a positive statistic may indicate either upregulation in females and downregulation in males or a higher increase or a lower decrease of the variable in human females with AUD. On the other hand, a negative statistic may indicate either upregulation in males and downregulation in females or a higher increase or a lower decrease of the variable in human males with AUD. In this comparison, the behavior of each lipid across the groups must be assessed a posteriori, examining female (IF) and male (IM) comparisons (Fig. S2).

### Web platform

All data and results generated in the different steps of bioinformatics strategy analysis are available on a web platform (https://bioinfo.cipf.es/sal-chronics), which is freely accessible to any user and allows the confirmation of the results described in this manuscript. The front-end was developed using the Angular Framework, the interactive graphics used in this web resource have been implemented with plotly [30], and the exploratory analysis cluster plot was generated with the ggplot2 R package [31].

This easy-to-use resource is divided into seven sections: 1) a summary of analysis results; the detailed results of the 2) class annotation results; 3) exploratory analysis; 4) differential abundance between experimental groups; 5) LSEA results; and 6) metabolic lipid network results, where the user can interact with the web platform through graphics and tables and search for specific information related to lipid species or classes; and 7-8), which include methods, bioinformatics scripts, and supplementary material.

## Results

### Sex-based differences in lipid species and class lipid profiling of plasma EVs isolated from individuals with AUD

The lipidomic profile of plasma EVs from females and males with AUD and control individuals revealed 311 and 264 lipid compounds by using the negative and positive ion modes, respectively. After normalizing the sample data, lipid species were labeled by their ion mode. RefMet and LIPID MAPS databases were used to classify all lipids (575 species) in different subclasses and their upper levels (super and main classes) (Fig. 3A and Table S3). The descriptive analysis of the lipid composition showed an enrichment of TAG, PC and SM subclasses in plasma EVs (Fig. 3A). Regarding the lipid abundance distribution of the different subclasses (Fig. 3B), all patient groups displayed similar abundance profiles, except for OxPC-O and CAR. However, hierarchical clustering of the EV lipid species, regardless of their subclass, revealed that the lipid profiles of the four experimental groups were distinct from one another, as shown in Figure 3C-D. The samples were separated by disease (individuals with AUD vs. controls) and by sex (Fig. 3C). Moreover, part of the variance was explained by the sex (PC1) and by the disease (PC2) (Fig. 3D).

**Figure 3.**
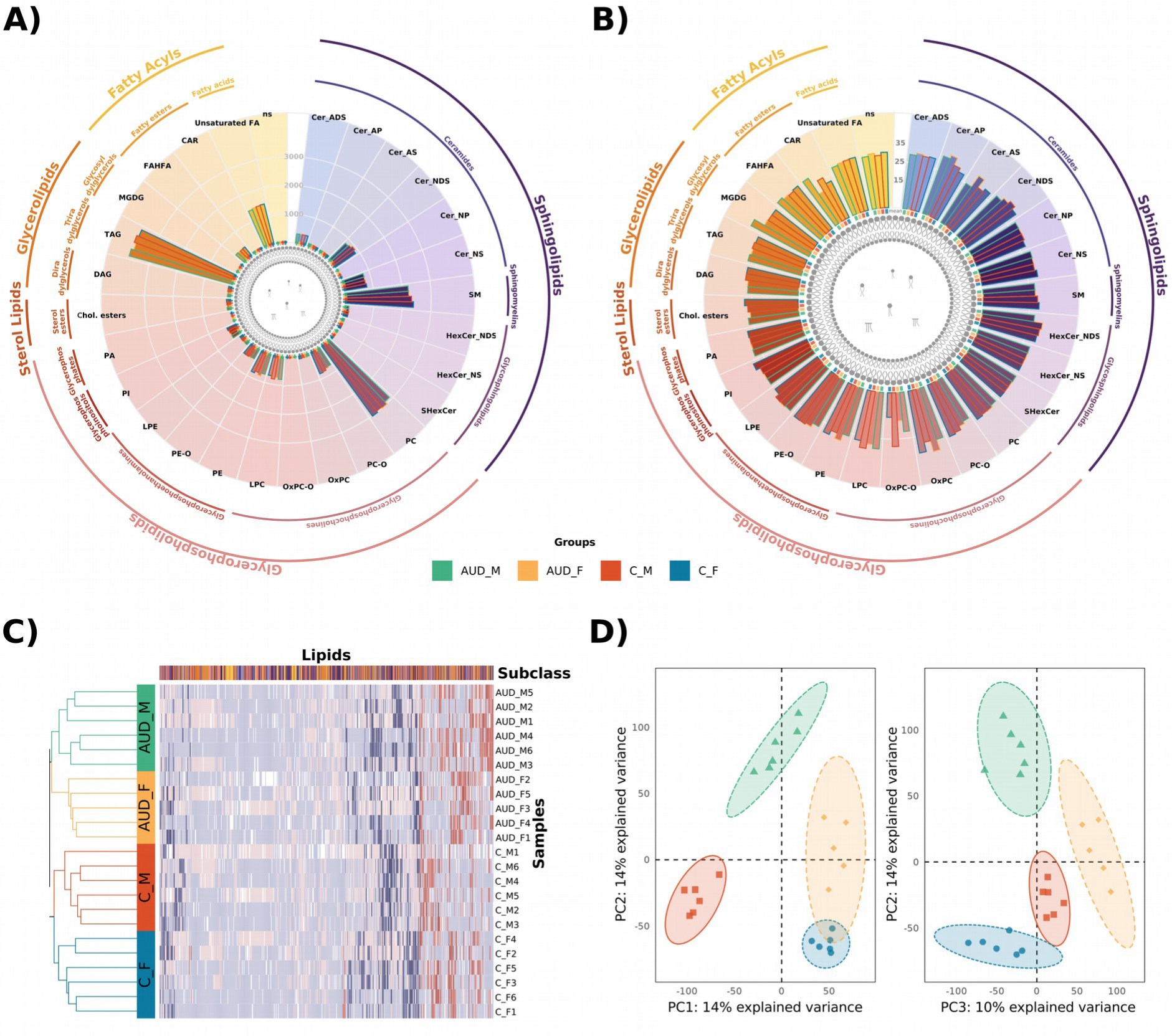
Overview of samples, individual groups and class lipids. **(A)** Sum of total abundance for each patient group for each subclass. The inner lines indicate the main class and the outer lines the super class. The color of the border of the bars indicates the individual group. **(B)** Median of abundance for each patient group for each subclass. The inner lines indicate the main class and the outer lines the super class. The color of the border of the bars indicates the individual group. **(C)** Heatmap demonstrating the abundance patterns between lipids (columns) and samples (rows), and the column colors indicate the subclass. **(D)** Principal component analysis (PCA) of the samples by the patient groups.

To assess significant variations in lipid abundance in human plasma EVs, three comparisons were conducted: 1) females with AUD vs. control females (IF), 2) males with AUD vs. control males (IM), and 3) IF vs. IM (named IS to identify the impact of the sex). As shown in Figure 4A, 32 and 39 lipid species were significant (p-value ≤ 0.05), when comparing females and males with AUD to control individuals, respectively. Additionally, the IS comparison revealed that 15 lipids displayed a significant difference, indicating a sex-specific response to AUD. If we delve into the lipid subclasses of Figure 4A, to which significant lipids belong, showed differences between all comparisons. Specifically, the Cer_AP and Cer_AS subclasses revealed significant lipids that were more abundant only in females with AUD, while LPC and PE subclasses displayed significant lipids that were less abundant in females with AUD. The FAHFA subclass exhibited significant lipids that were more abundant only in males with AUD, whereas the PE-O subclass had a significant lipid that was less abundant in males with AUD. It is worth noting that the lipid species belongs to the PE-O subclass (Fig. 4A label N) showed lower abundance in males with AUD and appeared significant with logFC > 0 in the IS comparison.

**Figure 4.**
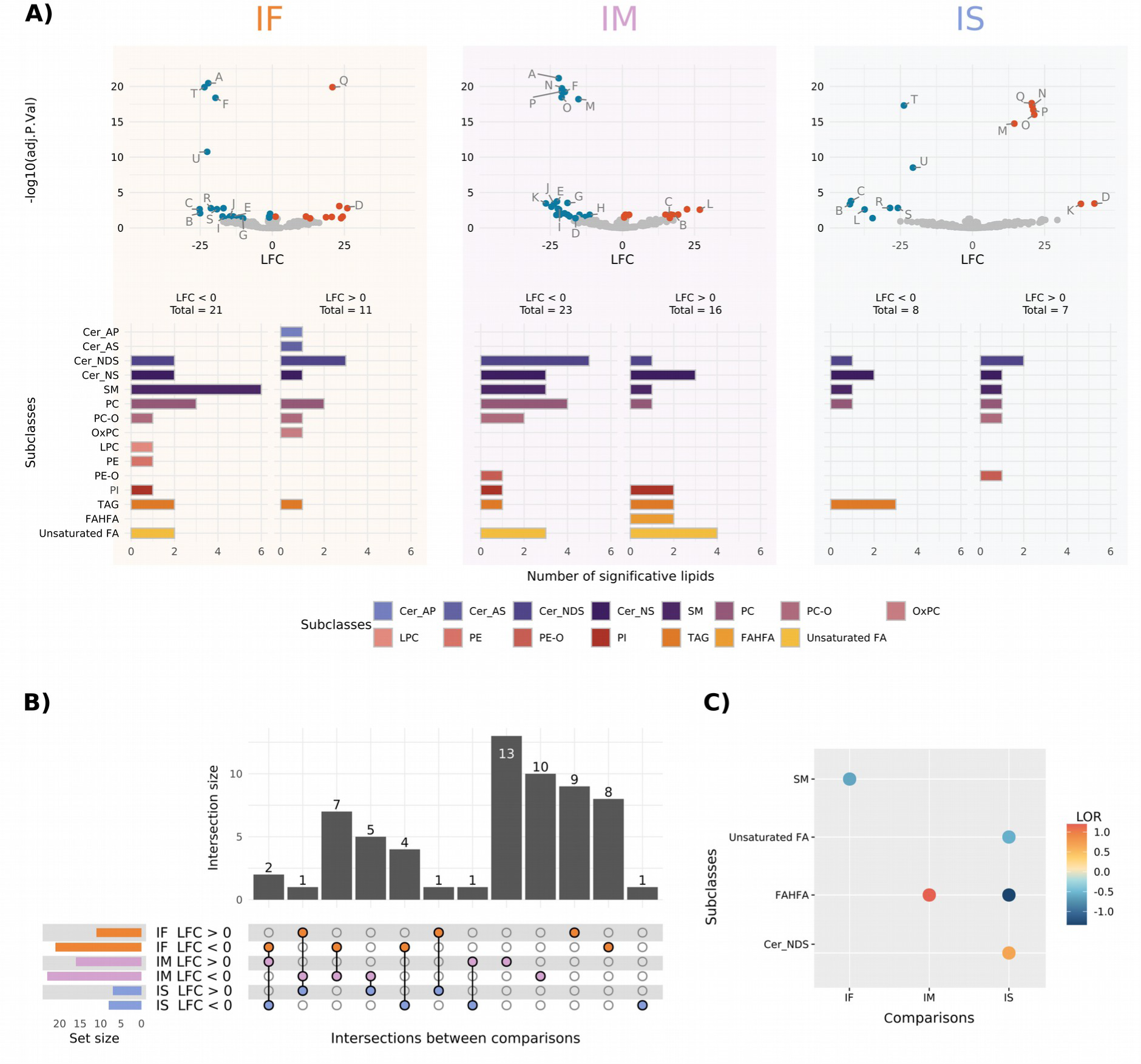
Summary of the differential abundance analysis and molecular lipid profile in each comparison. **(A)** Volcano plot summarizes all lipid data and bar plot displays significant lipids classified by subclass and LFC (logarithm of fold change). Over-abundant and under-abundant significant lipids are shown in red and blue, respectively (p-value adjusted ≤ 0.05). Non-significant lipids are represented in gray. Significant lipids in at least two comparisons are mentioned; A: Cer_NDS d39:1_neg, B: Cer_NDS d42:2 RT:12.673_neg, C: Cer_NS d18:1_22:0_neg, D: Cer_NS d18:1_24:1_neg, E: Cer_NS d18:2_23:0_neg, F: EtherPC 16:0e_18:2_neg, G: FA 22:0 RT:6.523_neg, H: PC 32:3 RT:6.415_pos, I: PI 18:0_18:2_neg, J: SM d18:2_24:0_neg, K: Cer_NDS d42:1_neg, L: Cer_NS d18:1_24:0_neg, M: EtherPC 38:5e_neg, N: EtherPE 16:1e_22:6_neg, O: PC 18:2_20:4_neg, P: SM d37:2_pos, Q: Cer_NDS d18:0_18:0 RT:12.135_neg, R: PC 39:4_pos, S: SM d42:4_neg, T: TG 18:1_18:1_20:1_pos, U: TG 54:7_pos. **(B)** Upset plot of the differential abundance analysis. Data from each comparison are separated according to the LFC sign. Horizontal bars indicate the number of significant lipids in each comparison (a specific color for each comparison). Vertical bars indicate the lipids included in the intersection of the groups denoted with a colored dot underneath. A colored dot under a bar indicates the specificity of the lipids in this group. **(C)** Analysis of the enriched significant lipid subclasses by LSEA. The dot colors represent the sign and magnitude of the change (LOR). IF - impact of AUD in females (orange); IM - impact of AUD in males (purple); IS - impact of sex in AUD (blue).

Analyzing the number of significant lipids shared between the different comparisons, we identified sex-specific lipid species (Fig. 4B). Specifically, 22 lipid species were female specific, 29 species were male specific and only 10 lipid species were shared between the IF and IM comparisons. Considering these last 10 lipids species, 7 were less abundant in female and male individuals with AUD (see Table S4). On the other hand, the other 3 lipid species displayed an opposite abundance and were significant in the IS comparison (see Table S4). Furthermore, 15 lipids exhibited sex-based differences (IS comparison) in the AUD response. Some of them were also found to be significant in the IF and/or IM comparisons, while 1 lipid was exclusively significant in the IS comparison.

The LSEA results (Fig. 4C) displayed a significant enrichment of the SM subclass in IF comparison, being less represented in AUD than in control females (negative LOR value). IM comparison in the LSEA results suggested that the FAHFA subclass was more represented in AUD males than in control individuals (positive LOR value). We also observed a significantly higher enrichment of the Cer_NDS subclass in AUD females compared to AUD males. However, a significantly higher enrichment of Unsaturated FA and FAHFA subclasses in AUD males compared to AUD females (SI comparison) was also shown.

### Sex-based differences in the lipid network of plasma EVs isolated from individuals with AUD

LINEX^2^ aims to obtain a biological interpretation of lipidomics data. Figure 5 represents the global network of lipid species, which provides qualitative associations between species based on defined reaction types. The major part of the reactions observed were related to fatty acid modify/removal (orange and blue edges). If we delve into the study, Figure 6 depicted similar qualitative associations as Figure 5, while also providing quantitative information about alterations in lipid levels between AUD and control conditions in females (IF) (Fig. 6A) and males (IM) (Fig. 6B). Since the colored spherical node represents higher lipid abundance, the IF network (Fig. 6A) revealed some lipids with increased abundance (not significant) in both control females (blue nodes) and with AUD (red nodes), with a uniform distribution within the network. However, there were more abundant lipids in control males (Fig. 6B, blue nodes). Additionally, the lipids that exhibited higher abundance in males with AUD (larger spherical nodes) indicates statistical significance. The edge color in the network indicates the correlation change of reaction connecting two nodes. Figure 6A-B shows distinct patterns between the sexes, and some lipids exhibit opposite Log Fold Change (LFC) values. The network zoomed view, represented by the lipid species Cer(18:1;O2/24:1), Cer(18:1;O2/22:0), and Cer(42:2;O2) (Fig. 6C, F), denoted as Cer_NS d18:1_24:1_neg, Cer_NS d18:1_22:0_neg, and Cer_NDS d42:2 RT:12.673_neg (Fig. 4), also confirmed sex significant differences. The lipid DG(18:1/18:1) (highlight node Fig. 6D, G) showed significant connections with several other lipids, displaying different correlations in the IF and IM comparisons. For instance, its interaction with TG (18:1/18:1/21:0) shows a non-significant correlation in females with AUD, but a significant correlation in control females in the IF comparison. In the IM comparison, this correlation is significant in males with AUD, but not significant in control males. Furthermore, the global network also showed notable differences in the correlations between both sexes (Fig. 6E, H). In females, green edges indicate significant reactions in control individuals (not significant in females with AUD). In contrast, males showed an opposite network zoomed view (blue edges).

**Figure 5.**
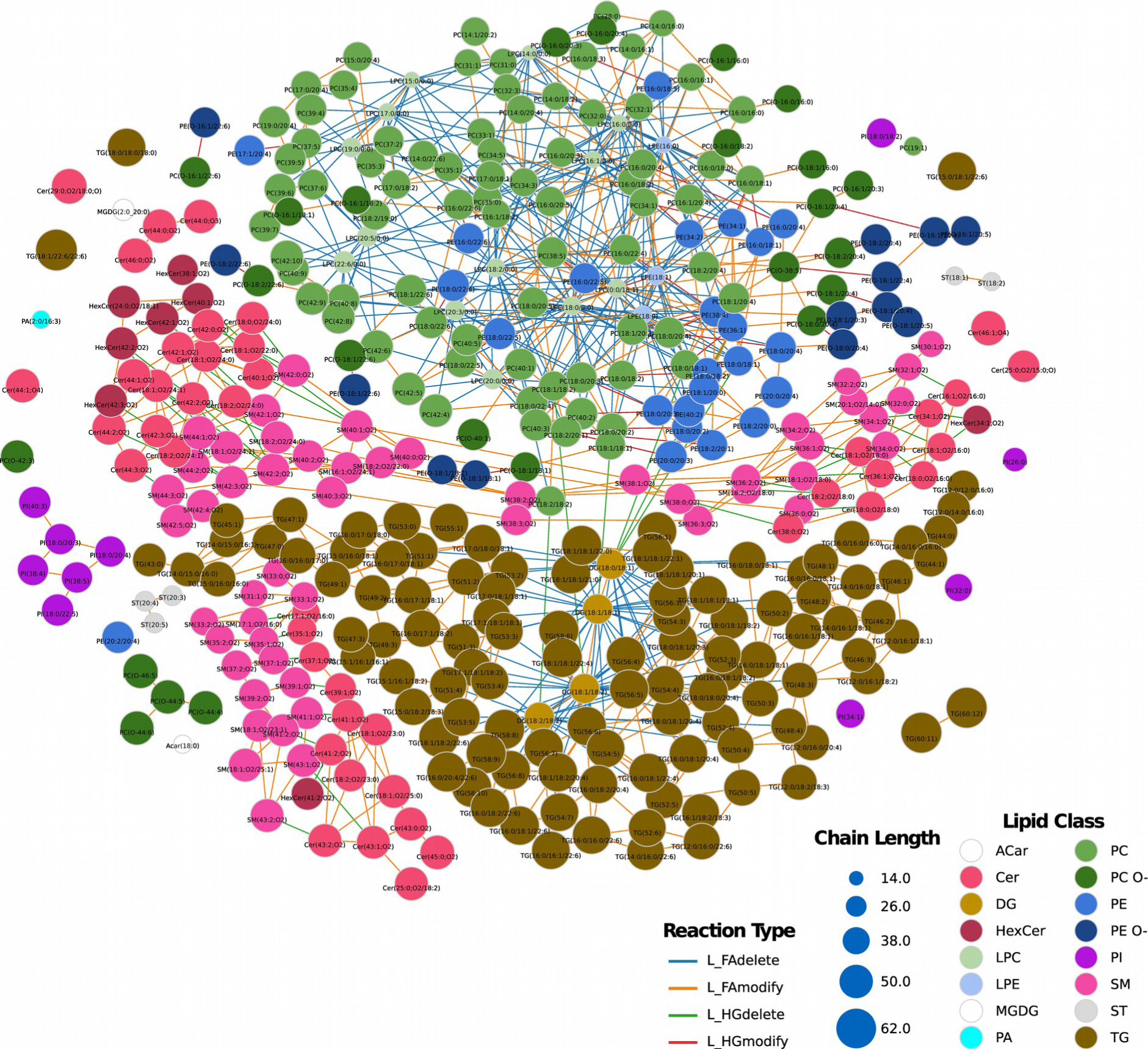
LINEX^2^ lipid network based on LC–MS/MS lipid data. Colored spherical nodes depict lipid classes. Edge colors indicate the type of reaction, which connects the nodes. For further exploration and analysis, an interactive version of the network, along with all other LINEX ^2^ analyses are accessible in an HTML file, through the link https://drive.google.com/file/d/1hOTFa4uZS8zfuU9LO_UhaNeadGiy4ayZ/view?usp=drive_link.

**Figure 6.**
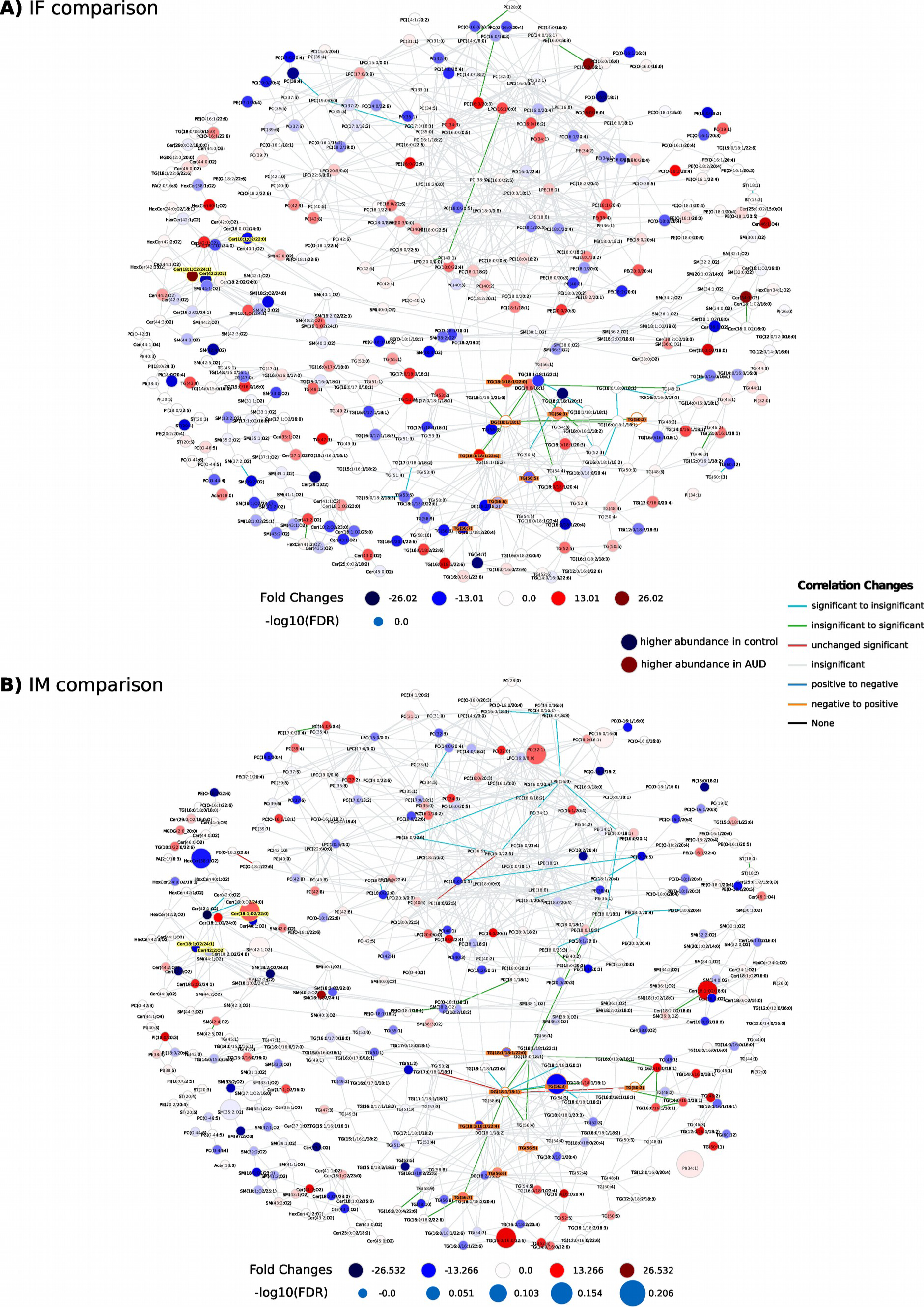
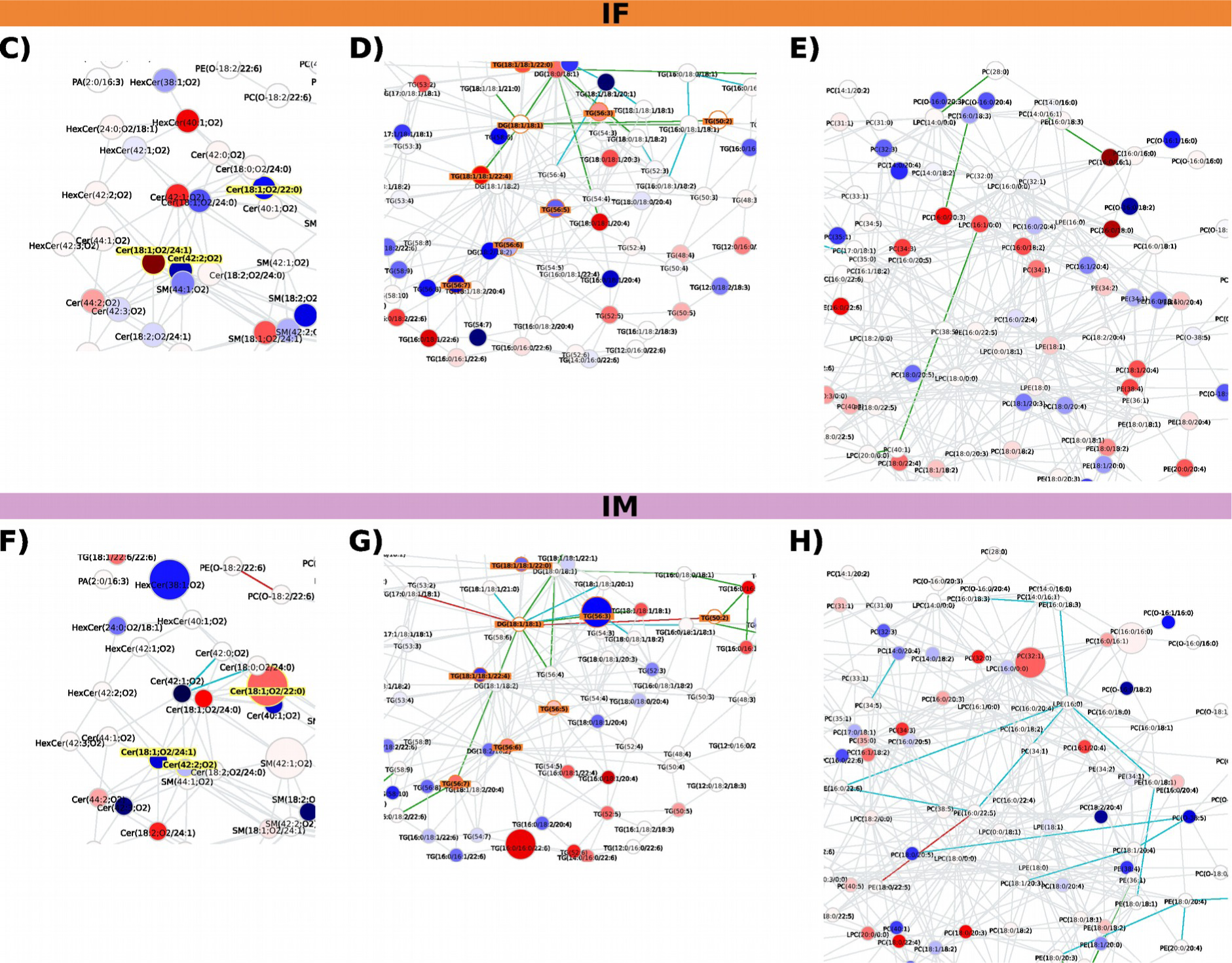
Lipidomics data visualized with LINEX^2^ for IF and IM comparisons and network zoomed views. IF **(A)** and IM **(B)** comparisons in the lipid network. Red spherical nodes represent lipids with a positive LFC from control to patients with AUD (higher abundance in AUD), whereas blue nodes indicate negative LFC (higher abundance in control). The spherical node sizes indicate the negative log10 FDR corrected p-values of lipid species between the AUD and control groups (a larger node size represents a higher level of statistical significance). Lipid network and other LINEX^2^ analyses can be explored in an interactive version, available as an HTML file at https://drive.google.com/file/d/1hOTFa4uZS8zfuU9LO_UhaNeadGiy4ayZ/view?usp=drive_link. **(C-H)** Network zoomed views of specific lipid species and reactions. The same three specific enzymatic reactions are observed in the IF network (C-E) and IM network (F-H). In the lipid network, red spherical nodes represent lipids with a positive LF, from control to AUD (higher abundance in AUD), and blue nodes indicate negative LFC (higher abundance in control). Edges are colored by changes of correlation for lipids from the patients with AUD to healthy individuals. For example, green indicates a non-statistically significant correlation in the AUD condition and a statistically significant correlation in the controls, where the correlation has the same sign.

### Sex-based differences in lipid enzymatic dysregulation of plasma EVs isolated from individuals with AUD

Using lipid class reactions from common metabolic databases through a network enrichment algorithm [7], we can determine enzymatic dysregulation from our EVs lipidomics data. Figure 7 shows the enrichment networks generated by LINEX^2^ based on the global networks (Fig. 5-6). Figure 7A highlights the dysregulated subnetworks that maximize the reaction difference between the AUD and control conditions in females and males. The resulting subnetworks include only PC and LPC lipid species in females, and Cer and SM in males, suggesting that the enzymatic dysregulations are involved in different types of biochemical reaction in both sexes, transforming the lipid species into each other.

**Figure 7.**
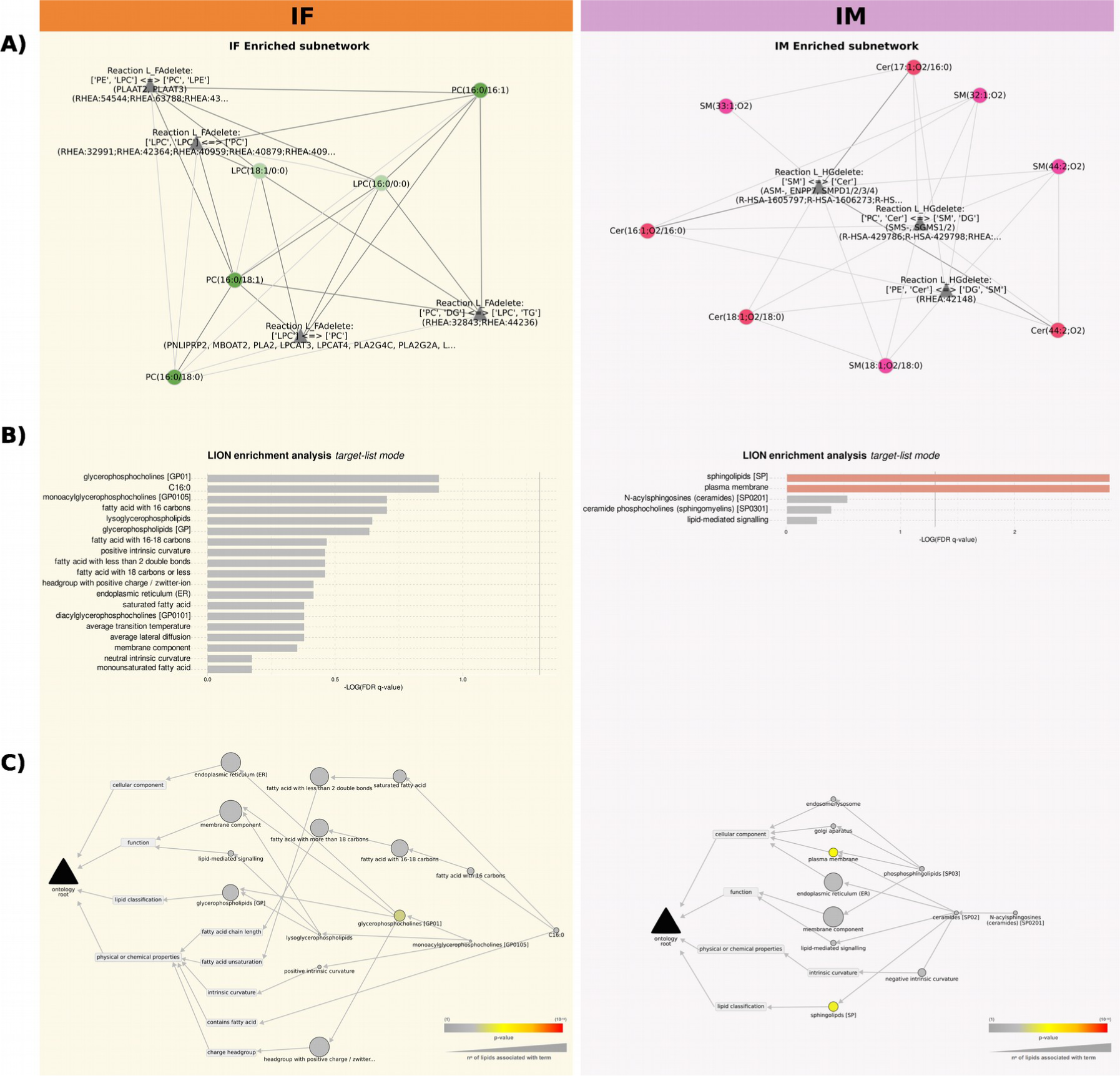
Enrichment networks generated by LINEX2 based on the global networks. **(A)** LINEX^2^ enrichment subnetworks for IF and IM comparisons. Spherical nodes represent lipid species and triangular nodes represent the reaction type. **(B)** The most enriched ontology terms result from using the lipids in the subnetworks as targets in the target list mode. **(C)** A hierarchical network displaying the most enriched ontology terms result from using the lipids in the subnetworks as targets in the target list mode. The node colors are scaled by the raw p-value, and the node size indicates the number of lipids involved in each ontology sterm.

Some differences appeared between sexes in Lipid Ontology (LION) enrichment analysis, using the lipids in the subnetwork as targets in the target list mode (Fig. 7B). Notably, females exhibited ontology terms related to membrane activity and stability, such as “positive intrinsic curvature”, “headgroup with positive charge/zwitter-ion”, “lipid-mediated signaling” or “endoplasmic reticulum”. These concepts are usually associated with the lipid class glycerophosphocholine, and the terms related to this class were enriched. In males, the two most significant terms were “sphingolipids” and “plasma membrane”, which are related to cell membrane and lipid signaling pathways. It was noted that females showed ontology terms related with “positive intrinsic curvature” of the membrane, while male presented “negative intrinsic curvature” terms (Fig. 7C), both of them related to EV biogenesis. Thus, the properties of the lipidome assigned by the LION algorithm could suggest alterations of the lipids involved in membrane remodeling and lipid-mediated signaling in the EVs from patients with AUD in a different pattern between sexes.

### Web platform

The web platform (https://bioinfo.cipf.es/sal-chronics) contains detailed information regarding the complementary computational approaches involved in this study. This resource includes statistical indicators of each performed analysis, which users can explore to identify their own profiles of interest. This open resource hopes to contribute to data sharing between researchers, the elaboration of innovative studies, and the discovery of new findings.

## Discussion

Preclinical studies have highlighted the importance of considering a better understanding of the biological and metabolic pathways involved in AUD to the development of new medications and diagnostic methods to treat AUD. In this sense, most studies have focused on the utility of miRNAs and proteins carried by EVs as plasma biomarkers. However, our results demonstrated for the first time that in females with AUD, LPC and PC lipids, along with enzymes like phospholipases and acyltransferases, show substantial changes associated with cancer progression and neuroinflammation. Moreover, males with AUD exhibited dysregulation of Cer and SM lipid species, involving sphingomyelinases, sphingomyelin phosphodiesterase, and sphingomyelin synthase, potentially contributing to ethanol-induced hepatotoxicity. Additionally, computational analysis highlights sex-specific variations in lipid properties within EVs associated with vesicle fusion processes.

Considering that EVs can be derived from all body cells and circulate in body fluids, the characterization of their lipids could give us information associated with the cell or tissue of origin and their functional states [32]. The distribution of the different lipid species in absolute amounts demonstrated that the most abundant lipid subclasses identified were PC, SM and TAG. Whereas PC is one of the most abundant lipid classes in EVs derived by neural cells [33], SM is involved in the biogenesis of the EVs, as well as one of the most abundant classes in brain-derived EVs [34, 35]. For instance, SM d18:2_24:0 is a novel lipid specie, which could be used as a potential biomarker in females and males with AUD. However, the presence of TAG in EVs could be derived from a secretory autophagy pathway [36]. In addition, TAG could also be transferred from lipoproteins to exosomes once released into the bloodstream [37], suggesting the absence of lipoprotein contamination in EVs isolation procedures [5].

We have previously reported that acute ethanol intoxication induced a higher enrichment of distinct plasma EV lipid species (e.g., LPC, PA, FAHFA) in human female adolescents than in males. These lipid classes participate in the formation, release, and uptake of EVs and the activation of the immune response [5]. Following the same sex differential analysis procedure in patients with AUD, our results reported that the subclasses LPC and PE were less abundant in females with AUD than in healthy individuals. LPC is enriched in EVs related to proinflammatory functions and also participates in EV biogenesis [38]. Moreover, LPC promotes demyelination, through the activation of inflammatory responses in CNS and by inducing pyroptosis in microglia [39]. Indeed, alcohol-induced proinflammatory molecules in the periphery may provoke neuroinflammation by crossing the brain blood barrier [40]. A general decline in plasmalogen lipids, mainly PC and PE subclasses, has been described in multiple brain regions in Alzheimer’s disease [33], which could be associated with an increase of the oxidative stress, the inflammatory responses and the neuronal cell death [41, 42]. However, other studies reported high and low levels of PC and PE in highly metastatic breast cancer, respectively [43]. In addition, our results also showed that most of the ceramide lipid species (e.g. Cer_NS d18:1_24:1, Cer_NS d18:1_22:0, Cer_NDS d42:2 RT:12.673) exhibited sex-specific abundance with an opposite pattern between the sexes. The subclasses Cer_AP and Cer_AS were more abundant in females with AUD, whereas some lipids belonging to Cer_NS and Cer_NDS were less abundant. This increase in Cer_AS species, along with a decrease in Cer_NS and Cer_NDS, has been described in a mouse model of metachromatic leukodystrophy, suggesting that alpha-hydroxylation of ceramides may play a role in the brain pathology of this disease (e.g., demyelination and motor dysfunction) [44].

Another important lipid class is the fatty acids, which have been associated with inflammation [45] and neurotransmitter release [46], through cell surface and intracellular receptors, modifying membrane composition, influencing cell signaling, gene expression, and lipid mediator production [45]. Our results show that unsaturated FA (main class Fatty acids) has a negative LOR in the IS comparison, indicating a class enrichment in males with AUD compared to female patients. FAs have been implicated in neural cell pathology in lysosomal storage diseases, including Metachromatic Leukodystrophy, a condition characterized by lipid accumulation in the brain, spinal cord, and peripheral nerves [44]. Furthermore, although the FAHFA subclass emerges as a significant and more abundant/enriched lipid in males with AUD, except for its role as anti-inflammatory function, little is known about the involvement of the FAHFA in the biological processes [47].

The incorporation of LINEX^2^ lipid network enrichment to our data also provides the basis for a knowledge-driven integration of lipidomics with proteomics data, by connecting enzymatic activity to lipid species [7]. The resulting network analysis showed greater substrate-product changes in females with AUD for the lipid reactions of the LPC and PC subclasses, which include the enzymes phospholipases and acyltransferases (e.g., LPCAT3/4). Enzymes, such as LPCAT1, have been reported to be upregulated in human colorectal adenocarcinoma [48] and in human metastatic prostate cancer [49], suggesting an involvement of the LPC metabolism with the cancer progression. In addition, the PLA2-activated neuroinflammatory pathways through the upregulation of the oxidative stress status is induced by binge alcohol treatment in adult rats and in organotypic hippocampal-entorhinal cortical slice cultures [50]. Our results also demonstrated that the enzyme PLA2G2A is upregulated in females with AUD. This enzyme, which shows lysophospholipase, transacylase and PLA2 activities [51], has antimicrobial function, by degrading bacterial membrane and by releasing pro-inflammatory eicosanoids from inflammatory cell-derived EVs [52].

Our results also showed an enzymatic dysregulation of Cer and SM lipid species in males with AUD. Previous studies have reported alterations in the levels of various sphingolipids, including Cer and SM, in human chronic alcohol-related liver disease [53] and in individuals with high alcohol consumption [54]. Furthermore, the enzymes involved in these substrate-product reactions are the sphingomyelinases (e.g., ASM, ENPP7, SMPD family, and SGMS1), which have been associated with chronic alcohol consumption [55]. In this line, recent studies have proved the activation of the sphingomyelinase activity in ethanol-treated microglial cells [56] and high sphingomyelinase protein levels in the pathogenesis of alcoholic liver disease [57]. Considering that the enzymes involved in the sphingolipid metabolism, might play a role to mediate the hepatotoxic effects of ethanol [58], the activation of ASMase and generation of C16-ceramide could sensitize hepatocytes to the effects of TNF-α [59]. According to our results, a sex-based difference study reported high levels of the serum ASMase activity in alcohol-dependent male patients [60].

Taking into account that lipids exhibit a plethora of structural and signaling functions, it is important to consider not only the biosynthesis of lipids, but also the change in biophysical properties. Consequently, we performed a comprehensive computational analysis of the lipidome, using both network-based and lipid property-related methods through the LION algorithm, to evaluate membrane remodeling and lipid-mediated signaling in EVs. Interestingly, our results showed a LION-term enrichment featuring “positive intrinsic curvature” in females with AUD, whereas in males with AUD exhibited a “negative intrinsic curvature”. Lipids with positive intrinsic curvature, such as LPC, have been found to hinder stalk formation during vesicle fusion [61], facilitating the expansion of fusion pores [62]. However, whereas lipids with greater negative curvature, such as PE and DAG, are critical players in the fusion, lipids of lesser negative curvature, including phosphatidic acid, are more likely to play a modulatory role [63]. The vesicle fusion processes are significantly influenced by lipids with negative curvature [63, 64], such as oleic acid or DAG, which tends to promote stalk formation and inhibit pore expansion [65]. It is important to note that the formation and expansion of fusion pores in SNARE-dependent vesicle fusion are essential steps for neurotransmitter release and vesicle recycling during exocytosis [66].

Our interest in this study was to provide data from individual lipid species, which is essential for a rigorous lipidomic pathway analysis. Indeed, lipid species of the same lipid class can behave completely differently, leading to distinct biological functions. However, certain limitations can also be presented in the analysis. For instance, 1) the lack of both standardization of lipid nomenclature and its integration into the different computational tools used (e.g., while FAHFA is significant in lipid abundance, it could not be included in the LINEX2 software), 2) lipid databases, such as LIPID MAPS and HMDB, contain general information of lipid class biology, and 3) LINEX2 software details lipid species and their enzymatic activity, although its use does not allow a full control and provides aleatory results based on the algorithm. Regarding the EVs used, these microvesicles have similar size and protein markers to exosomes, although it is currently impossible to specifically identify them as exosomes.

In conclusion, this study employed an innovative strategy based on a network enrichment algorithm to gain insights into the sex-specific dysregulation of lipid enzymatic reactions in patients with AUD. Notably, our findings unveiled sex-based differences in lipid profiles related to EVs biogenesis and may underlie inflammatory and neurodegenerative responses. These methodological advancements have deepened our understanding of lipid metabolism and its regulatory mechanisms, facilitating the identification of novel lipid targets and the potential discovery of sex-specific clinical biomarkers for AUD.

## Declarations

## Ethics approval and consent to participate

Human plasma samples were used in accordance with the Declaration of Helsinki and was approved by the Ethics Committee of the University Hospital of Salamanca (March, 2010), and written informed consent was obtained from each participant.

## Consent for publication

Not applicable.

## Availability of data and materials

The datasets generated and analyzed during the current study, as well as programming scripts are available in the Zenodo repository, https://doi.org/…, and in a web platform: https://bioinfo.cipf.es/sal-chronics.

## Authors’ contributions

CPC analyzed the data; MP and FGG designed and supervised the bioinformatics analysis; PC and MM obtained human plasma samples; SM isolated EVs from human plasma; CPC designed and implemented the web tool; CPC, MP, FGG and CGR wrote the manuscript; CPC designed the graphical abstract; CPC, MP, FGG and CGR helped in the interpretation of the results; CPC, CGR, MM, CG, MP and FGG writing-review and editing; MP and FGG conceived the work. All authors read and approved the final manuscript.

## Acknowledgments

The authors thank the Principe Felipe Research Center (CIPF) for providing access to the cluster, which is co-funded by European Regional Development Funds (FEDER) in Valencian Community 2014-2020. The authors also thank the Genomics and Proteomics Unit at the University of Alicante, the Electron Microscopy Service at the Príncipe Felipe Research Centre, Irene Soler-Sáez for designing Figure S2, and Stuart P. Atkinson for reviewing the manuscript.

## Abbreviations

ASM: Acid Sphingomyelinase
ENPP7: Ectonucleotide pyrophosphatase/phosphodiesterase family member 7
LPCAT 1/3/4: Lysophosphatidylcholine Acyltransferases 1/3/4
PLA2: Phospholipase A2
SMPD: Sphingomyelin Phosphodiesterase
SGMS1: Sphingomyelin Synthase 1

